# Quantifying phenotypic variability and fitness in finite microbial populations

**DOI:** 10.1101/680066

**Authors:** Ethan Levien, Jane Kondev, Ariel Amir

## Abstract

In isogenic microbial populations, phenotypic variability is generated by a combination of intrinsic factors, specified by cell physiology, and environmental factors. Here we address the question: how does phenotypic variability of a microbial population affect its fitness? While this question has previously been studied for exponentially growing populations, the situation when the population size is kept fixed has received much less attention. We show that in competition experiments with multiple microbial species, the fitness of the population can be determined from the distribution of phenotypes, provided all variability is due to intrinsic factors. We then explore how robust the relationship between fitness and phenotypic variability is to environmental fluctuations. We find that this relationship breaks down in the presence of environmental fluctuations, and derive a simple formula relating the average fitness of a population to the phenotype distribution and fluctuations in the instantaneous population growth rate. Using published experimental data we demonstrate how our formulas can be used to discriminate between intrinsic and environmental contributions to phenotypic diversity.

## Introduction

A central problem in microbiology is understanding how measurable phenotypic traits such as a cell’s size, growth rate, or expression of a gene affect population dynamics [1, 2, 3, 4]. This problem is made difficult by the fact that there is rarely a well-defined mapping from genotype to phenotype, instead, a single genotype can give rise to a distribution of phenotypic traits throughout a population’s history. This distribution can be generated at the cellular level due to *intrinsic* factors, such as the stochasticity of biochemical reactions [5, 6, 7, 8, 9, 10], or due to *environmental* fluctuations that affect the entire population [11, 12, 13, 14, 15]. It is now well established that a population’s long term growth rate can not be determined by averaging single-cell traits, such as the growth rates or generation times of cells, but is instead determined by the variability and heritability of phenotypes measured throughout a population. The discrepancy is an example of *non-ergodicity* and results from epigenetic heritability of phenotypic traits that is present in all populations. It is therefore necessary to quantify the distribution of a phenotypic trait, and not just the average, in order to deduce how phenotypes are affect fitness.

Measurements of phenotypic variability are commonly obtained in microfluidic devices, such as the *dynamics cytometer* pictured in Figure 1 (A), where a fixed number of cells are grown for long periods of time in controlled conditions [16, 17, 18, 19]. Such experimental setups allow one to track single-cell dynamics over hundreds of generations, a task that would be impossible if the population continued to grow exponentially. By tracking cells in a finite population over many generations, detailed information about the variability and epigenetic heritability of phenotypic traits can be obtained. How does one relate this information to some measure of the population’s fitness? Many previous studies have explored how population growth is related to the single-cell dynamics of an exponentially growing population in a constant environment [1, 20, 21, 22, 23, 19, 24, 25]. In the setting of exponential growth, a proxy for fitness is the population growth rate Λ, defined by the relation *N* ∼ e^Λ*t*^, where *N* is the number of cells at time *t*. Most notably, it was shown that the population growth rate, Λ, can be computed from the distribution of single-cell generation times taken over the entire history of the population using the *Euler-Lotka equation*, shown in Figure 1 (B). The Euler-Lotka equation establishes a link between the distribution of phenotypes and the fitness of the population, but it generally requires knowledge of the entire *lineage tree* depicted in Figure 1 (B). In the finite population, one cannot obtain the entire lineage tree because most of the cells are expelled from the population. Instead, one must analyze data from a “pruned” lineage tree such as the one depicted in Figure 1 (B). The objective of our study is to understand how the fitness of a finite population can be quantified from this, more limited, lineage data.

**Figure 1:**
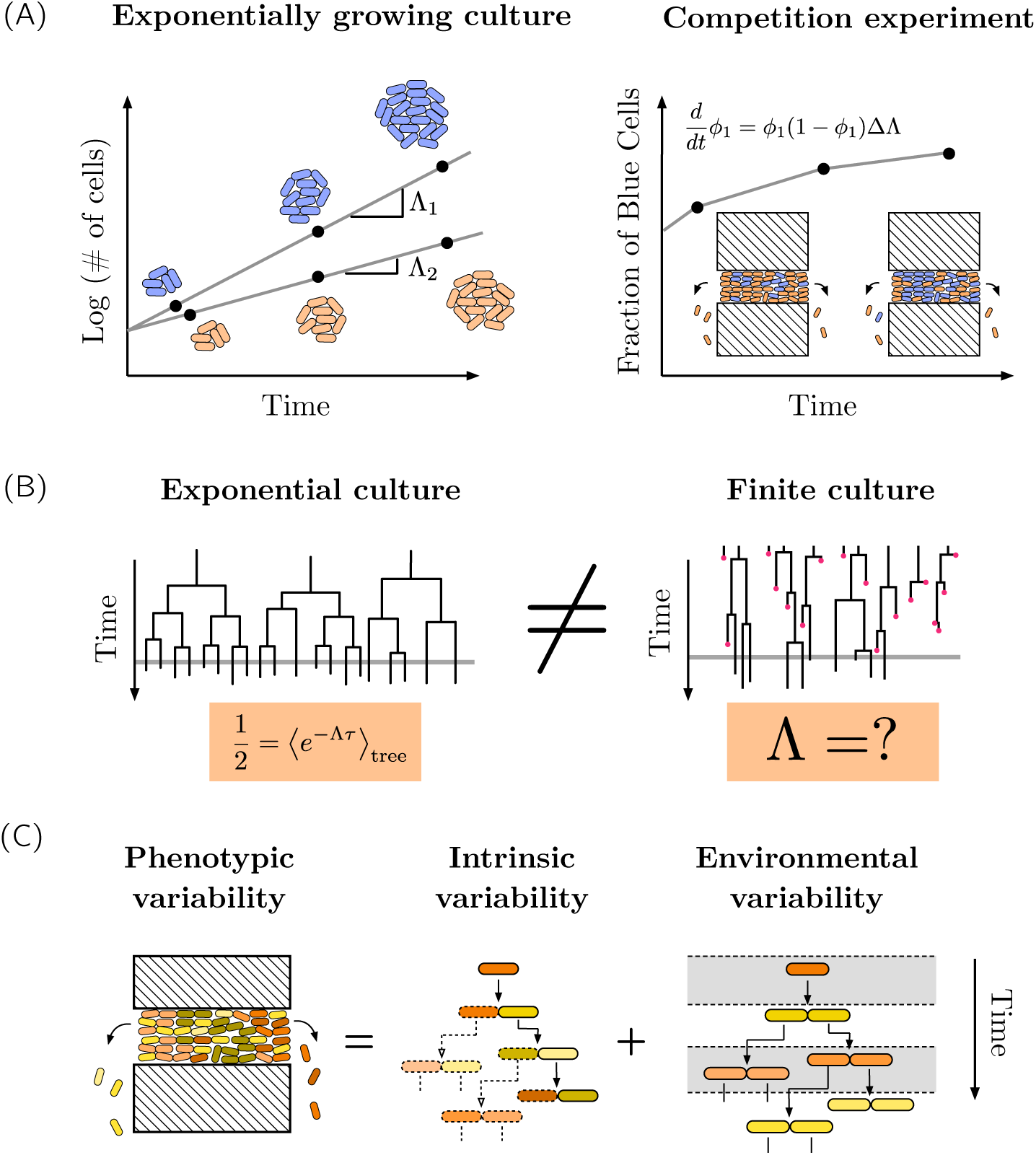
Population dynamics, lineage data and sources of phenotypic variability. (A) (left) In a population growing exponentially the population growth rate, Λ, a proxy for fitness, can be computed from temporal dynamics of the number of cells. (right) In a finite culture containing two species, the fraction of one species, *ϕ*_1_, typically evolves according to the logistic growth equation with rate parameter ΔΛ representing the difference in growth rates of the individual species. (B) Lineage data obtained form the different experiments in (A). Each vertical line represents a cell and the length of the line its generation time. In the exponentially growing populations, the lineage data can be related to the population growth rate, Λ, via the Euler-Lotka equation (orange box). The average ⟨ · ⟩_tree_ refers to the average over every branch in the tree. In finite culture experiments it is unclear how to obtain the distribution of all cells on the tree because most of the cells are expelled from the population. (C) (left) A diagram of a device used to collect single cell lineage data. (middle) A model of cell proliferation with only intrinsic variability. In such a model, the generation times along a lineage obey a Markov process. (right) Temporal fluctuations in the environment (gray and white areas) may also induce phenotypic variability.

By studying the relationship between the statistical properties of lineage trees in exponentially growing and finite cultures, we derive an analog of the Euler-Lotka equation that requires only knowledge of the pruned lineage data. Just as one can compute the population growth rate in an exponentially growing culture by measuring the slope of ln *N* (*t*), in the finite culture the population growth rate can be estimated directly by observing the rate at which cells are expelled from the culture, or the *dilution rate*. Importantly, the accuracy of these estimates depends on whether the phenotypic variability comes from intrinsic or environmental factors. The former arises due to stochasticity at the cellular level that causes different cells throughout the population to have distinct phenotypic traits, while the later is a result of environmental fluctuations affecting unrelated cells simultaneously; see Figure 1 (C). We find that in the presence of environmental fluctuations, the population growth rate cannot be determined solely from the phenotype distribution, and we derive a formula for the deviations of the population growth rate from the prediction of the Euler-Lotka equation in this setting.

## Results

### General model of intrinsic variability

We start by introducing a general modeling framework allowing us to study intrinsic variability in an arbitrary (and possibly multi-variate) phenotypic trait *x*. We assume each cell is born with a value of *x* drawn from a distribution *f* (*x* |*x*′) depending on the phenotype *x*′ of the mother cell. For example, *x* might represent the initial concentration of a protein which is expressed stochastically over the course of a cell cycle, or some macroscopic phenotype, such as the growth rate. After a time *τ* (*x*), the cell divides and produces two new cells with generation times drawn from the same distribution. This framework can capture any possible form of *intrinsic* (as opposed to environmental) generation time variability through the appropriate choices of the phenotype *x* and distribution *f*. Therefore, existing models can be derived as specific cases of this framework. These include models where division is asymmetric [26, 27, 28, 25] and where the relationship between successive generation times is non-monotonic and nonlinear, such as the kicked cell-cycle model used to describe circadian rhythms [29].

As a concrete example, we will often refer to a specific model of cell-size regulation, which is described in detail in Methods and [1]. The central assumptions of this model are that single cells increase in volume exponentially with time and divide (up to some noise) at a specified volume. The strategy for selecting the volume at division is known as the *cell-size regulation strategy*. In the notation of our modeling framework, the phenotype 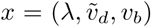 where *λ, v*_*d*_ and *v*_*b*_ are the cell’s growth rate, volume at division and volume at birth respectively. The transition operator *f* (*x*| *x*′) is then dependent on the specific cell-size regulation strategy (see Methods).

### Growth rate analysis in the finite culture

In the setting where a population described by our model grows exponentially, the problem of determining the population growth rate Λ from *f* (*x* |*x*′) has been explored in a number of recent studies for specific choices of the observable *x* [30, 20, 21]. In our general modeling framework, Λ, along with the distribution of phenotypes among all cells in the exponentially growing population tree, *ψ*_tree_(*x*), obeys the recursive equation (see Methods)

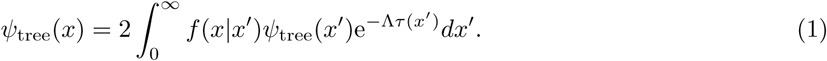

Integrating with respect to *x* retrieves the Euler–Lotka formula [31],

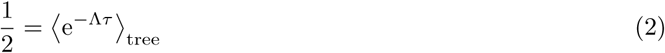

Equations 1 and 2 determine the unknowns *ψ*_tree_(*x*) and Λ in terms of the model of phenotypic variability, which is encoded in *f* (*x*|*x*′). Given that *f* (*x*|*x*′) is sufficiently smooth in *x* and *x*′, uniqueness of these solutions can be established using the theory of integral equations [31], and we assume throughout that these conditions are satisfied. As discussed in the introduction, only in the simple case where phenotypes are uncorrelated across generations (*f* (*x*|*x*′) = *f* (*x*)) is this distribution equal to the distribution of phenotypes along a lineage, which we refer to as the *lineage distribution* and denote by *ψ*_lin_(*x*); see Figure 3 (A). Hence while the lineage distribution may be the most natural distribution to consider because it does not require us to grow the population exponentially, it is generally less informative if we are interested in the population growth.

**Figure 2:**
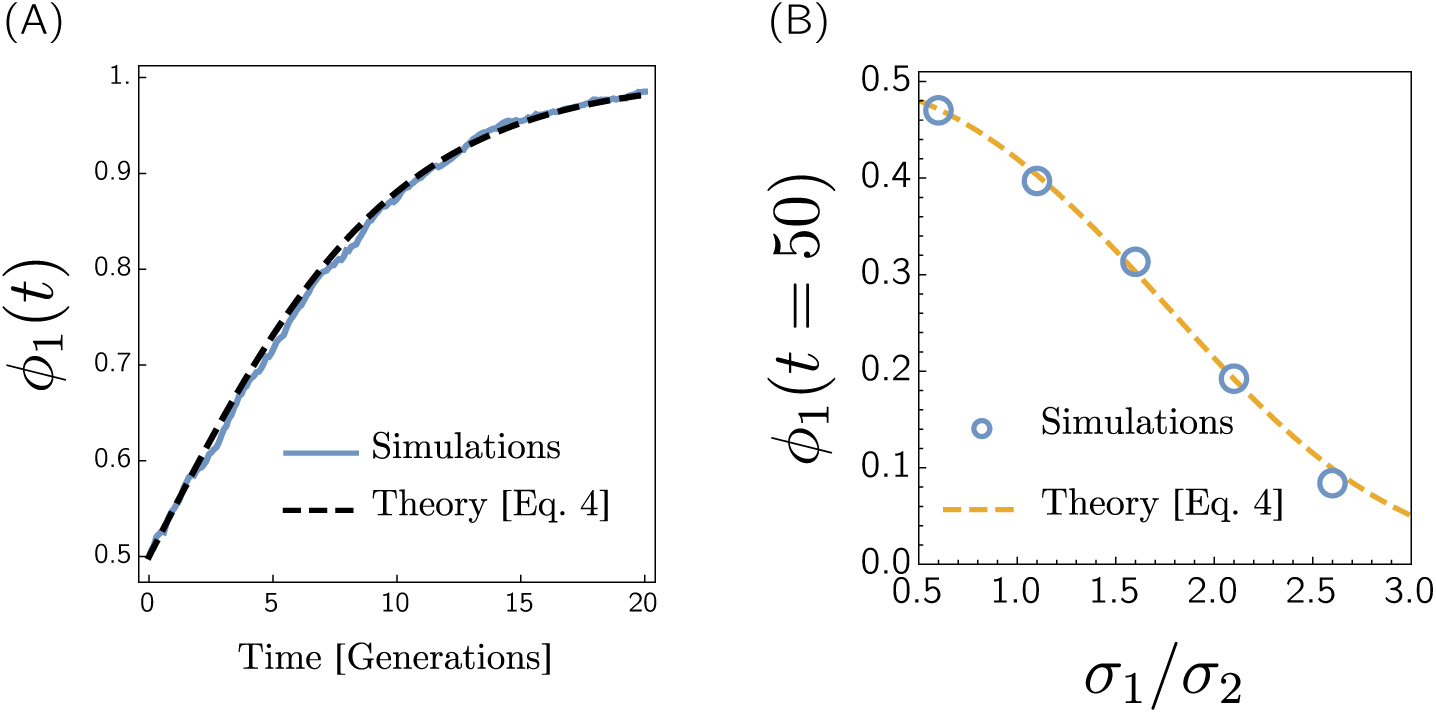
Population growth predicts fitness in the finite culture. (A) Sample path of *ϕ*_1_(*t*) starting with a culture equal parts species 1 and 2. Each population is modeled using the cell-size regulation model. We have non-dimensioqnalized by taking ⟨*λ* ⟩_lin_ = 1 and define a generation as the doubling time of a cell with the average growth rate, hence 1 generation = ln(2) units of time. (B) *ϕ*_1_(*t*) at *t* = 20 generations for two populations with the same average growth rate and different values of 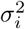, the variance in growth rates. We have fixed 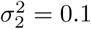 and increased *σ*_1_. For both species growth rates are uncorrelated across generations. Each circle represents the average of 1000 simulations for the same parameter values.

**Figure 3:**
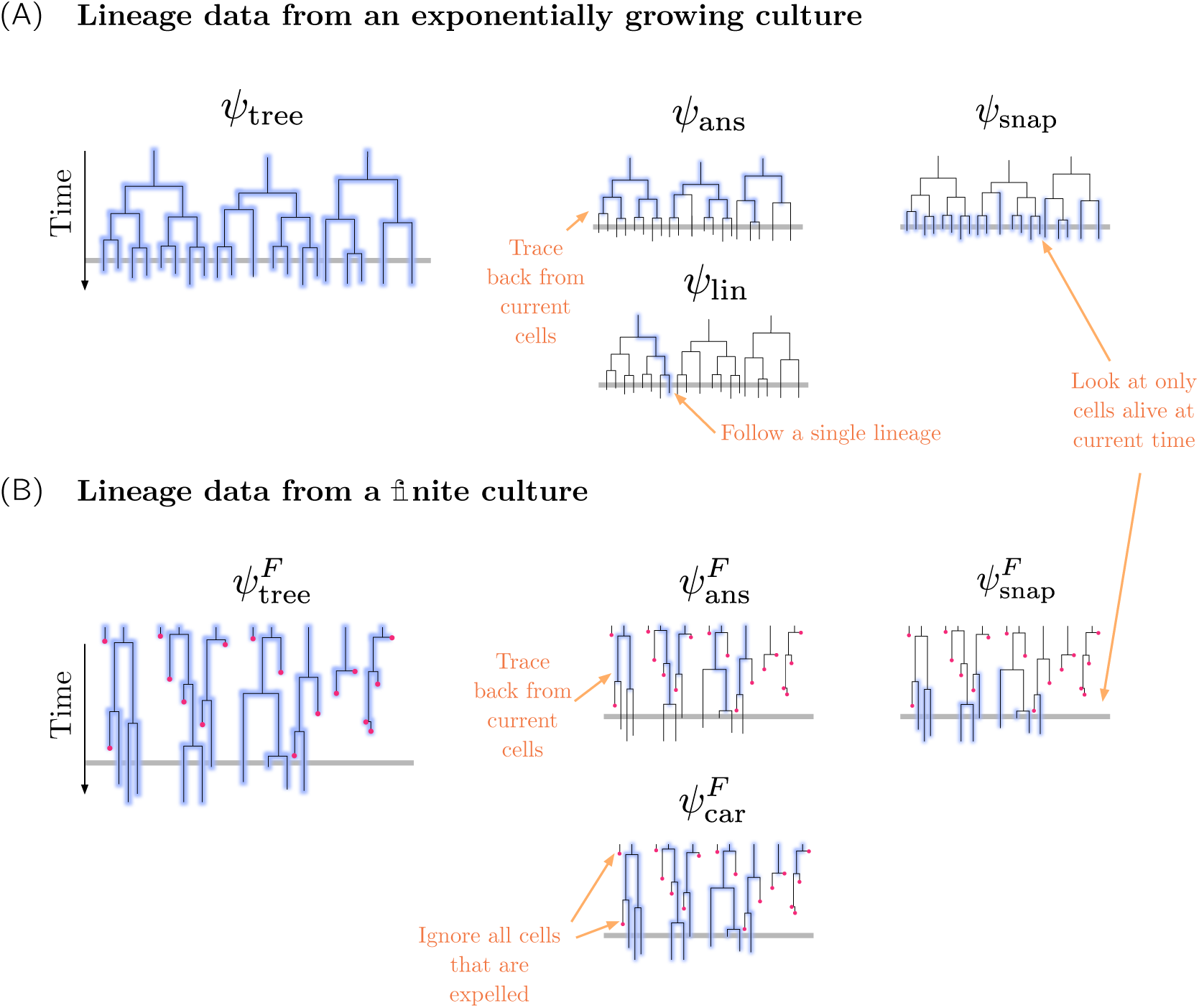
Methods of sampling lineage trees. (A) (left) The various ways of sampling phenotypes from a lineage tree obtained in an exponential culture. The gray horizontal line on the right panel indicates the current time of observation. The blue highlighted lines correspond to cells that are sampled to obtain the distribution indicated above the lineage tree. For example, to obtain *ψ*_ans_ we sample all the cell in the tree excluding those that are currently alive. (B) The same image for a finite culture. The red dots represent cells that are expelled from the culture before they are able to divide.

We now consider a model where *N*, the number of cells in the population, is fixed. In order to achieve this, every division event must correspond to a cell being expelled from the culture, which is approximately the case in a microfluidic device such as the one pictured in Figure 1 (B). In a simulation, this can be implemented by picking a random cell from the population at each division event and removing it from the culture. The resulting process is known as a *Moran process* in the mathematics literature [32]. Since the population is not growing, we can no longer obtain Λ from the exponential growth curves (given by the equation in Figure 1 (A)). Instead, one can measure growth in terms of the rate per cell at which cells are expelled from the population, or the *dilution rate* Λ_*D*_. Intuitively, if *N* is sufficiently large, the continuous expulsion of cells from the culture should have no influence on the relative frequencies of the phenotypes, and therefore the instantaneous rate at which cell volume is accumulated should be equal to Λ. On the other hand, since the size of the culture remains fixed, this must be balanced by the rate at which cells are expelled, implying Λ = Λ_*D*_. It follows that Λ_*D*_ should satisfy the same relation as Λ, namely,

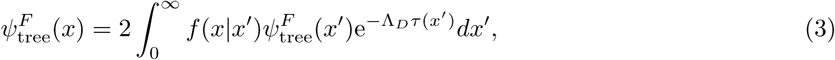

where we are now using 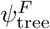 to represent the distribution of phenotypes over all cells throughout the history of the culture; see Figure 3 (B). As discussed in Methods, 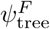 can also be interpreted as the distribution of phenotypes over all cells in a snapshot that have just been born. In general we will use the superscript *F* to indicate a distribution in the finite culture, but we omit this when there is no ambiguity. ^1^ For a rigorous derivation of Equation 3 we refer to Methods. When phenotypes are not correlated across generations, *f* (*x* | *x*′) = *f* (*x*), hence it follows from Equation 3 that 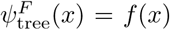. We may integrate with respect to *x* and replace Λ_*D*_ with Λ in order to obtain Equation 2. This relation was previously obtained Powell in a model without correlations across generations. For this reason, his derivation [31] does not clarify which distribution the average should be taken with respect to.

### Phenotype distribution predicts selective advantage in the finite culture

Having established that the Euler–Lotka equation can be generalized to the setting of the a finite culture, we now consider two species competing in a finite culture and show that the population growth rate is also a measure of fitness in this setting. We will once again assume that the number of cells (*N*) is sufficiently large so that any effects of order 1*/N* may be neglected. Before considering these competition dynamics directly, suppose that both species are grown separately in environments with unlimited space and nutrients, see Figure 1 (A). Under these conditions, the number *N*_*i*_ of cells in species *i* will then grow exponentially at some rate Λ_*i*_, hence 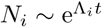. A simple calculation reveals that the fraction of species 1, *ϕ*_1_ = *N*_1_*/*(*N*_1_ +*N*_2_), evolves according to the logistic growth equation

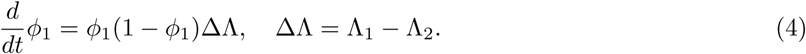

The quantity *S* = ΔΛ*/*Λ_2_ is known as the *selective advantage* of species 1 over species 2 and the sign of the quantity determines which species eventually takes over the culture. We now imagine placing both species in a finite culture that can hold exactly *N* cells; see the left panel in Figure 1 (B). As before, this is implemented by removing a randomly selected cell from the culture every time a cell division occurs. We will assume that nutrients are not limiting growth so that the rate at which cells of a given species divide does not depend on the relative abundances of each species. Heuristically, we can obtain the evolution of the frequency *ϕ*_1_(*t*) from the exponentially growing population by uniformly sampling *N* cells from the exponentially growing population at each time, therefore it is plausible to assume that Equation 4 describes the evolution of the frequency *ϕ*_1_ in the finite culture as well. A systematic derivation of Equation 4 for two phenotypically heterogeneous species along with the generalization to the case of *M* species is given in Methods and the agreement between the logistic growth Equation 4 and numerical simulations of two species competing in a finite culture is shown in Figure 2.

In the context of the cell-size regulation model, previous work has shown that if the mean growth rate is fixed along a lineage, variability in single-cell growth rates decreases the population growth rate for biologically realistic parameter values [20]. Therefore, it follows from our analysis that in a finite culture variability in growth rates will reduce a species’ competitive advantage. This is shown numerically in Figure 2, where we have simulated the competition between two species with different levels of growth-rate variability, but identical mean growth rates.

### The relationship between methods of sampling phenotype data

The tree distribution, which is required to compute the population growth rate using Equation 2, is only one of a number of ways to quantify phenotypic variability in a lineage tree, yet it is not always the most convenient to work with. For example, in the device pictured in Figure 1 (A) it is difficult to track the generation times of cells that are expelled from the culture, yet these are needed in order to compute 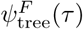. Therefore, it is useful to understand the relationship between the different distributions that can be obtained from the lineage tree. A related question is how distributions of phenotype over subsets of a lineage tree differ in exponential and finite cultures. In some cases the exponential and finite cultures are equivalent. For example, consider the current cells distribution, *ψ*_snap_(*x*). This is the distribution of phenotypes in a snapshot of the population at a given time; see Figure 3. One can show that in both finite and exponentially growing populations taking a snapshot of phenotypes yields the same distribution (see Methods). However, as we will show below this equivalence of distributions does not hold for phenotypes sampled over the ancestors of the current cells in the population.

Another important distribution is given by the expression 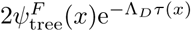; this term is twice the integrand of the Euler-Lotka equation, which implies it must be a probability distribution, as it is normalized. As was previously noted by Powell [31] in the context of a model with uncorrelated generation times, this can be interpreted as the distribution of *x* over all cells that divide before being expelled from the culture. Following Powell’s terminology, we define the *carrier distribution* as

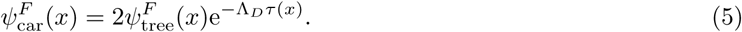

This relationship is verified numerically for the size-control model in Figure S.1. The subset of the lineage tree which gives this distribution is depicted in Figure 3 (B). To derive Powell’s interpretation of the carrier distribution in the more general setting, first note that 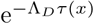 is the probability that a cell stays in the culture until it divides, given that is has phenotype *x*. Also, the probability that any cell throughout the history of the population divides in the culture must be 1*/*2 in order for the number of cells to remain fixed. The interpretation of 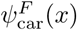 then follows from an application of Bayes’ Theorem where 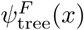 and 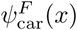 play the role of the *prior* and *posterior*, respectively. In the exponentially growing culture the distribution 2*ψ*_*tree*_(*x*)e^−Λτ(*x*)^ has a different interpretation; it is equal to the distribution over all the current cell’s ancestors, *ψ*_ans_(*x*) (this has been referred to as the *branch distribution* in [20]). This distribution is obtained by sampling all cells in the tree excluding those that are currently alive, as shown in Figure 3 (B). The fact that *ψ*_car_(*x*) = *ψ*_ans_(*x*) suggests there may be an alternative interpretation of these distributions that is common to both types of cultures. In fact, in both the continuous culture and exponentially growing culture, the carrier (or branch) distribution can be interpreted as the distribution of phenotypes of cells that have just divided, or the *division distribution ψ*_div_(*x*) (see Methods for a detailed discussion of this distribution).

In practice the distribution 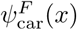 may be easier to obtain than 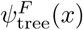 because sampling 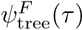 requires knowledge of generation times of cells that are expelled from the culture. In a microfluidic device these cells may be difficult to track, and additionally, their growth may be not be statistically identically to those cells that remain in the culture due to the different conditions in and out of the microfluidic chamber where cells are tracked. This motivates us to develop a method for estimating the population growth rate, Λ, from the available lineage data. To this end, we utilize the normalization of 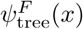 to obtain

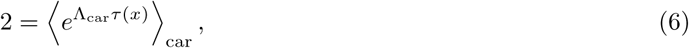

where we will denote the estimate of Λ obtained from this equation as Λ_car_ in order to distinguish it from the dilution rate estimate Λ_*D*_.

#### The ancestral distribution

In the exponential culture the ancestral distribution can be interpreted as the distribution over all the current cell’s ancestors. How might the distribution over the ancestral cells in the finite culture be related to *ψ*_ans_(*x*) in the exponential culture (or equivalently, *ψ*_car_(*x*))? Because all cells have a common ancestor we can compute *ψ*_ans_(*x*) by taking only the ancestor cells of this common ancestor; as we trace back in time far enough the more recent cells make up a negligible fraction of the observed phenotypes. *ψ*_ans_(*x*) is therefore the distribution of phenotypes obtained by looking *backwards* along a lineage beginning from an arbitrary cell in the culture. If phenotypes are not correlated across generations, then the phenotypes sampled along the lineage are independent and this is therefore equivalent to sampling cells in the population that have divided in the culture; as we saw before, this is *ψ*_car_(*x*). However, if there are correlations across generations, we need to consider how the ancestor’s phenotypes along a lineage are biased by the decedent’s phenotypes. This is captured by the recursive formula for *ψ*_ans_(*x*) (derived in Methods):

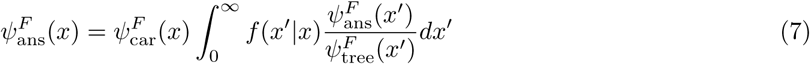

From this equation, it is easy to see that. 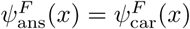 precisely when *f* (*x*|*x*′) = *f* (*x*), as expected from the argument above.

Our results concerning the relationship between phenotype distributions in exponential and finite cultures are summarized in Figure 4 (A) and (B). To explicitly demonstrate these relationships we performed simulations of the cell-size regulation model (discussed in Methods). Results of these simulations are shown in Figure 4 (C).

**Figure 4:**
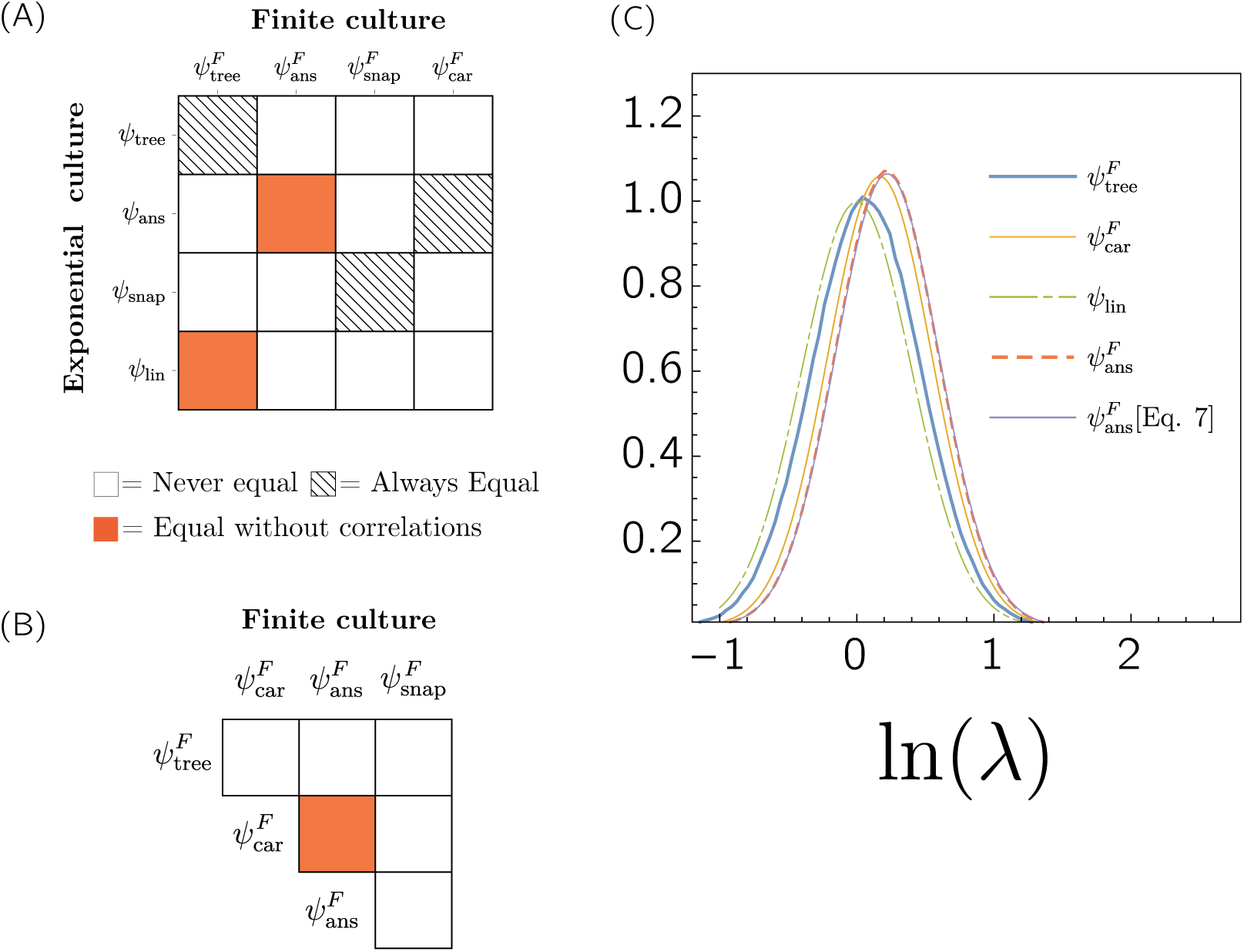
The relationship between various methods for sampling lineage data. (A) The relationship between distributions in the finite and exponentially growing culture. (B) The relationship between different ways of sampling the phenotype distributions in the finite culture. An expanded table that includes the birth and death distributions is shown in Figure S.3. (C) Plots of the various distributions of single-cell growth rates obtained from simulations of the cell-size regulation model with 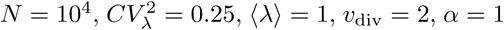 and *ρ*_*λ*_ = 0.2. The lineage distribution, *λ* which can be computed analytically for this model, is also shown. For the ancestral distribution, we have plotted both the histogram of ln *λ* over all ancestral cells (red dashed line) as well as evaluated the right hand side of Equation 7 using the carrier and tree distributions (purple line).

### Environmental fluctuations limit the predictive power of the phenotype distribution

In the previous section we derived a relationship between the population growth rate and the lineage data under the assumption that there is only intrinsic variability of phenotypes in the populations. However, it is very rarely the case that this assumption is satisfied in natural biological settings, and even in highly controlled experiments it is difficult keep the environmental conditions constant. How do both environmental fluctuations and intrinsic variability interact to shape a population’s growth rate? While the Euler-Lotka equation is no longer satisfied if the phenotype distribution is affected by environmental fluctuations, we will show that the generation time statistics can still be related to the population growth rate provided one also has information about the fluctuations in the instantaneous population growth rate.

To be concrete, we first consider an extension of the cell-size regulation model in which cells grow at a rate *λ*[*S*_e_] that depends on a time-dependent external nutrient concentration, denoted *S*_e_(*t*). A natural choice for the growth-rate function is Monod’s law [33],

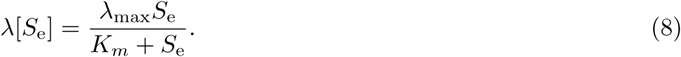

Here is *λ*_max_ is the maximum growth rate of a cell and *K*_*m*_ is the concentration at which a cell grows at half its maximum growth rate. As discussed in Methods, intrinsic variability is introduced by taking *λ*_max_ to be drawn from a normal distribution when a cell is born (with mean ⟨*λ* ⟩ and variance 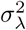) while environmental fluctuations come from perturbations of *S*_e_(*t*) around an average value *S*_0_. To this end, we define the environmental perturbation *ξ*(*t*) by *S*_e_(*t*) = *S*_0_(1 + *ξ*(*t*)). We will assume that ⟨*ξ*(*t*) ⟩ = 0 so that ⟨*S*_e_(*t*) ⟩_*T*_ = *S*_0_ and use 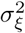to denote the strength of environmental fluctuations; 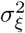 is given by the time average of *ξ*(*t*)^2^ over a long interval. In our simulations, we take *ξ*(*t*) to be an Ornstein–Uhlenbeck process; that is, *ξ*(*t*) undergoes a continuous time random walk in a quadratic potential with minimum at the origin. This process is characterized by a diffusion coefficient *D*_*ξ*_ and a relaxation time-scale *t*_*ξ*_. Importantly, *ξ*(*t*) is a *global* source of variability affecting all cells equally, while any variation in *λ*_max_ is due to intrinsic variability in the cell; see Figure 1 (C). Recall that the assumption of the cell-size regulation model is that a cell divides upon reaching a (possibly random) size _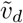_; therefore, the generation time *τ* of a cell born at time *t* satisfies 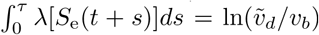. Adding environmental fluctuations to the cell-size regulation model in this way does not significantly affect the relationship between Λ_D_ and Λ. This can be seen numerically in Figure 5 (B) and an analytical justification is given in Methods. In contrast, the estimate Λ_car_ is a decreasing, linear function of 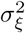 The specific from of this linear relationship depends on the parameters of Monod’s law according to

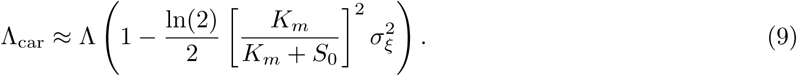

**Figure 5:**
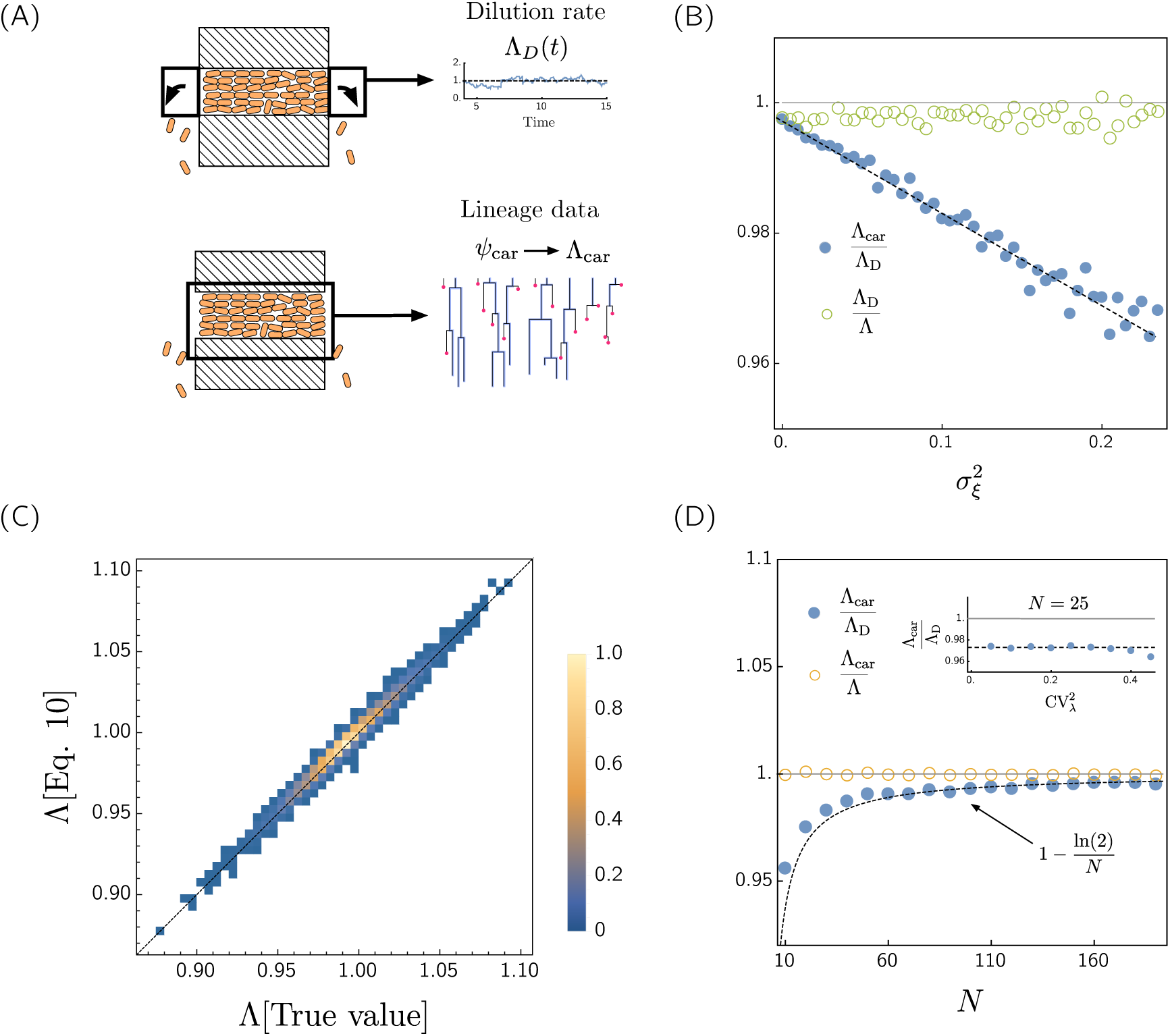
The relationship between growth rate predictions. (A) A summery of the different types of data and growth rate predictions that can theoretically be obtained from a finite culture. (top) The dilution rate is obtained by measuring the rate at which cells are expelled from the culture. (bottom) Single-cell tracking data can be used to obtain the carrier distribution (as well as other distribution of phenotypes). (B) Ratio of different estimates for the population growth rate, obtained from a finite population (Λ_car_, Λ_*D*_), and the population growth rate of an exponentially growing population (Λ), as a function of the environmental fluctuations. In all simulations, the instantaneous dilution rate is measured by sampling the number of cell expelled from the culture over intervals of length Δ*t* = 0.2 ⟨*τ* ⟩ and dividing by *N* Δ*t*. Here the environmental perturbation *ξ*(*t*) evolves according to an Ornstein-Uhlenbeck (OU) process. In this case the best estimate is clearly Λ_*D*_, which predicts the true population growth rate independently of the environmental fluctuations. Parameters are *N* = 500, *CV*_*λ*_ = 0.1, *ρ*_*λ*_ = 0 and *α* = 1. (C) The true population growth rate compared to the population growth rate obtain from Equation 10. The hue of each point represents the number of simulations with those measured growth rates (lighter points indicate more simulations). In order to compute the true value of Λ we have looked at the instantaneous change in volume over a small time interval *dt* = 0.1. Simulations were performed for the cell-size regulation model with *N* = 25, CV_*λ*_ = 0.2, *ρ*_*λ*_ = 0.1, *v*_*d*_ = 1, ⟨*λ* ⟩ = 1 and *α* = 1. The environment was simulated using a time-discretized OU process with relaxation timescale *t*_*ξ*_ = 10 and *D*_*ξ*_ was varied between 0 and 0.01. In Figure S.2, we use Equation 10 on the same synthetic data to compute 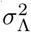 instead of Λ. (D) The same as (B), but showing the predictions for the different estimates of the population growth rate as a function of the population size, *N*. The parameter values used are the same as (B).

In Figure 5 (B) we have shown estimates of Λ_car_*/*Λ_*D*_ and Λ_*D*_*/*Λ from numerical simulations, which are consistent with our theoretical prediction.

Equation 9 implies that the population growth rate cannot be predicted solely from the distribution of generation times, instead, one needs both generation time statistics and information about the environmental fluctuations to predict the population growth rate. It is therefore natural to ask whether there exists a general relationship that connects the population growth rate to the generation time statistics via the environmental fluctuations. In the limit where the environment evolves on a time-scale longer than the typical generation time of a cell, such a relationship is given by (see Methods)

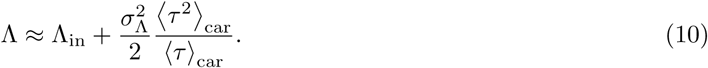

Here, Λ_in_ is the rate at which the population would grow if it had the same distribution of generation times in the carrier distribution, but no environmental variability, while 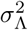 is the variance in Λ. As we have already shown, Λ_in_ is simply Λ_car_, and can therefore be computed from the carrier distribution. We have confirmed this relation numerically in Figure 5 (C). Equation 10 tells us that the phenotype distribution becomes less informative as environmental fluctuations increase. In Methods, we show that this relationship is also satisfied in a competition experiment between multiple species in a fluctuating environment, even if the environment has distinct effects on each species. In the next section we will discuss how 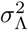 is related to the fluctuations in the dilution rate, which differ from the growth rate fluctuations in a small population.

### Population size effects

Because many experimental setups used to probe bacterial physiology are performed in devices that can only support between *N* = 15 and 40 cells, it is import to quantify the order 1*/N* terms in predictions of the population growth rate. The difference between the population growth rate, Λ, and the estimate obtained from the carrier distribution, Λ_car_, will be on the order of 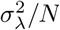. To see this, observe that in the limit where there is no phenotypic variability, Λ_car_ = Λ for all *N*. On the other hand, as 1*/N* approaches 0, Λ_car_ converges to Λ for all 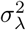 Therefore, we can express Λ_car_ as a Taylor expansion in 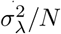 where the zeroth order term is Λ. This implies Λ_car_ gives a relatively accurate approximation to Λ. On the other hand, Λ_*D*_ deviates from Λ by a factor of order 1*/N*. Numerically, we have shown that

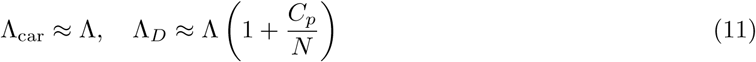

where *C*_*p*_ is a constant that is independent of the population size. For the cell-size control model, we have found *C*_*p*_ ≈ ln(2); see Figure 5 (D) and Figure S.4. The discrepancy between Λ_*D*_ and Λ arises because, in small populations with little phenotypic variability, all the cells tend to have similar ages and generation times. Therefore, cells divide at approximately the same time. As a result, the dilution rate tends to measure the inverse doubling time of the populations, 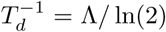. This can easily be seen in a population with no phenotypic variability, since in this case we will observe *N* cells expelled over a period *T*_*d*_, and therefore 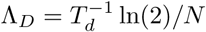.

Just as the average dilution rate depends on the populations size, the variance in growth rate, 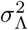, appearing in Equation 10 is distinct from the variance in the dilution rate, 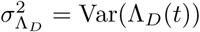. Formally, we can express the variance as a sum,

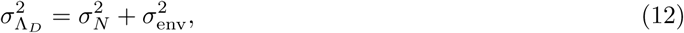

where 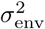 represents any fluctuations that do not vanish in an infinite population. In other words, for a given population, the term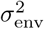 tells us how much the growth rate would fluctuate in a population with the same single-cell dynamics and environmental variability, but with an infinite number of cells. It follows immediately from this definition that 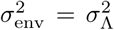 where 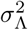 is the variance appearing in Equation 10. In many situations, it is desirable to determine 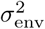 from data that is obtained in a small population where 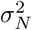 is non-negligible. In these situations, Equation 10 can be used to estimate 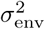 indirectly. To do so, one must have both the lineage data and a reliable estimate of the population growth rate. As we have shown, Equation 11 can be utilized to obtain such as estimate from Λ_*D*_. In the next section, we apply this procedure to experimental data.

### Application to experimental data

We now apply our results to analyze the experimental data original reported by Hashimoto *et al.* in [16]. In different experimental conditions, *Escherichia coli* were grown in a dynamics cytometer that can support approximately 10 to 50 cells. In [16], the population growth rate was estimated by solving Euler-Lotka equation for the tree distribution (Equation 2), by using the distribution of all cells that divide in the culture (the carrier distribution) weighted by their survival probability. This procedure gives an excellent approximation to the dilution rate, but is problematic in a small population where, as the authors observe, the dilution rate deviates from the population growth rate. The authors also computed the mother-daughter generation time correlations. In *E. coli*, generation times are typically slightly negatively correlated with its mother’s generation time (see [1] and Methods for an explanation in terms of the cell-size regulation model). However, it was found that in certain experiments there were relatively large positive generation time correlations; see Figure 6. One explanation for such correlations is that environmental fluctuations are influencing the growth statistics. We therefore expect that the environmental variability should be correlated with the mother-daughter generation time correlations.

**Figure 6:**
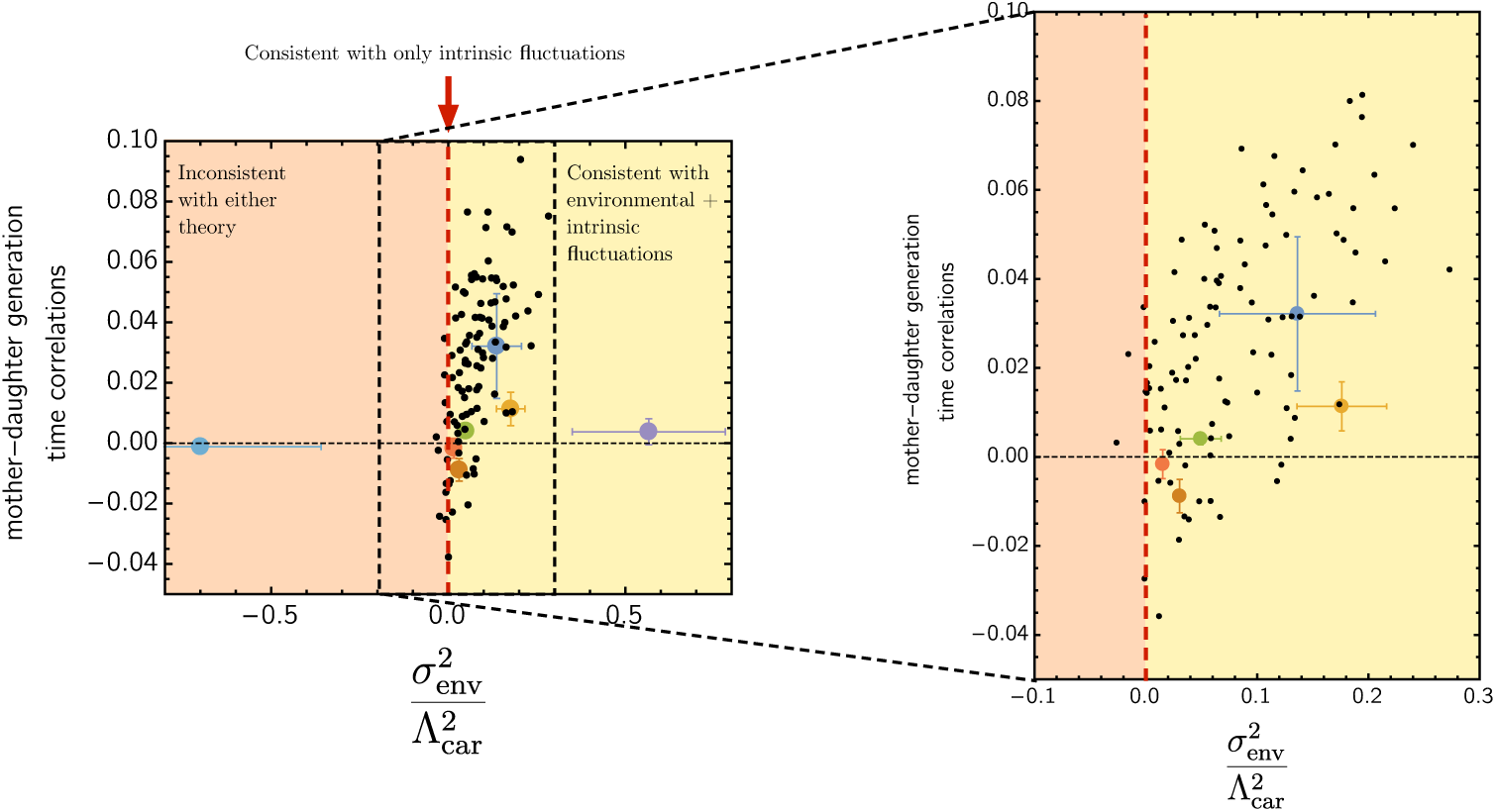
Predicting mother-daughter correlations in experimental data. The mother-daughter correlations as a function of 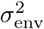 (computed by solving for 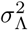 in Equation 10) for experimental data (colored points) and synthetic data (black points). The right panel shows a zoomed-in view of the region where most experimental data points lie. The synthetic data was generated by simulating the cell-size regulation model with 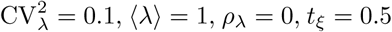, and various values of *D*_*ξ*_ (the diffusion coefficient of the OU process describing the environmental fluctuations) ranging from 0 to 0.1. Measurement error has been added by adding Gaussian noise to the growth rate and generation time measurements. Note that this leads to some data points with negative 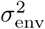, which is also observed in the experimental data. The fact that 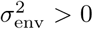 for many data points indicates that the phenotypic variability should not be interpreted as purely intrinsic. For additional information, see Figure S.3.

In order to test the predictive power of our theory, we used Equation 6 to directly solve for Λ_car_ and computed the dilution rate, Λ_*D*_, by averaging the instantaneous dilution rate over time. Using Equation 11, we are able to transform Λ_*D*_ into Λ, giving us all the ingredients to solve for 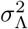 in Equation 10, which is equivalent to 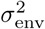 in Equation 12. We then plotted the mother-daughter correlations as a function of 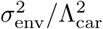; see Figure 6. This suggests that the positive generation time correlations are consistent with a model including both environmental and intrinsic noise, because all experiments with positive generation time correlations correspond to positive values of 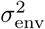. For one data point, 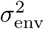 is negative, which is inconsistent with our theory, although this experiment is an extreme outlier in the data set. We have also compared the experimental data to synthetic data and observed that, after excluding outliers, both data sets obey the same trend. Note that the generation time correlations caused by environmental fluctuations may be canceled by cell-size regulation or other intrinsic effects in some cases, hence the fact that the correlations do not strictly increase with 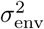 is consistent with our theory.

## Discussion

In this paper, we have studied the relationship between growth of phenotypically heterogeneous populations in finite and exponential cultures. While a number of studies have explored the dynamics of phenotypically heterogeneous populations under exponential growth conditions and shown how to relate lineage statistics to the population growth rate (e.g., [1, 20, 21, 22, 23, 19, 24]), precise measurements of phenotypic variability often need to be obtained in cultures containing a fixed number of cells. Because the growth process in not ergodic (the population averages are not equivalent to single-lineage averages) it is essential to distinguish between different ways of sampling phenotypes when analyzing data from these experiments. For example, an erroneous result is obtained if the lineage statistics are used to compute the population growth rate instead of the carrier distribution (the distribution taken over all cells that are not expelled from the culture). Moreover, in some situations one is limited in what distributions can be sampled. This the case in certain microfluidic devices where it is difficult to obtain the distribution of generation times of *all* cells because we do not have access to the generation times of cells that are expelled from the culture.

We have clarified the relationship between the different distributions in both the exponentially growing and finite culture, as well as derived a simple mapping between the various distributions in the two types of cultures. Importantly, distributions obtained in equivalent ways do not always have the same meaning in different types of cultures. In particular, the ancestral distribution in a finite culture is not equivalent to the ancestral distribution of the exponentially growing culture due to a bias towards surviving cells, yet both are obtained by looking at the history of all cells in the culture. A corollary of our mapping between the lineage statistics in different cultures is a formula for the population growth rate (given by Equation 6) that requires knowledge only of the lineage data obtained in a sufficiently large finite culture. In addition to relating the lineage statistics to the population growth rate, we have shown that the population growth rate predicts the fitness of a species in a competition experiment.

We have also considered how environmental fluctuations influence population growth predictions, and established a model-independent relationship between the fluctuations in the population growth rate and the growth rate prediction based on the phenotype distribution. Since single cell growth rate variability has been observed to change growth rate estimates by around several percent in *E. coli* [20], even small environmental fluctuations can conceal the intrinsic effects of phenotypic variability. We have established that the phenotype distribution becomes increasingly less informative as the environmental fluctuations in the population growth rate increase in amplitude (Equation 10). Hence, in the presence of environmental variability the fitness of a population is not determined solely by the distribution of phenotypes. We have tested the predictive power of the relationship between the population growth rate, the phenotype distribution and the environmental fluctuations on synthetic and experimental data. For most experimental data points, we see that there is a relationship between the environmental fluctuations predicted from Equation 10 and the mother-daughter generation time correlations. This is consistent with the observation that environmental fluctuations tend to create positive correlations between mother and daughter cell’s generation times.

Over the last decade there has been a growing interest in understanding the implications of phenotypic heterogeneity in the clinical context, yet understanding how to separate distinct sources of variability has proven to be a difficult problem [34, 11, 35]. For example, *Salmonella* grown in host macrophages exhibit increased phenotypic variability compared to *Salmonella* grown in *in vitro* cultures, yet the underlying causes of this variability, and in particular, whether it is induced by environmental fluctuations or stochastic gene expression is not known [11]. We have proposed one method for dissecting phenotypic variability in a growing population, and although we have motived this study by data obtained in microfluidic devices, one could in principle apply these same techniques to populations grown in more complex environments. Specifically, by tracking single cell dynamics of bacteria growth in macrophages, it may be possible to deduced whether the increased phenotypic variability was caused by environmental or intrinsic factors by examining the consistency of different growth rate predictions, as we have done here. This would require generalizing our framework to account for both spatial and temporal environmental fluctuations, which we have not included in our analysis, as much of the environmental variability is spatial in host environments [11]. Nonetheless, we believe this theory provides a promising starting point for future investigations of phenotypic variability.

In addition to data-driven applications of our theory, there are numerous theoretical questions concerning the fitness effects of phenotypic variability that we have left unanswered. For example, our analysis of the competition experiment has assumed that nutrients are not limiting growth and the dilution rate is adjusted to keep the population size fixed (such conditions can be created in a turbidostat [33]). We therefore do not account for chemostat-like conditions where species compete for limiting resources. In the future, it would be interesting to extend our analysis to account for such effects. It will particularly useful to explore how variability in different parameters effecting growth, such as maximal growth rate and yield, affect fitness. In addition, we have assumed that the competition occurs in a well-mixed environment, yet the geometry of the environment where the competition takes place can vastly influence the evolutionary dynamics [36]. It would be interesting to investigate the fitness effects of phenotypic variability in more complex environments where these factors play a role.

## Acknowledgements

We acknowledge Jie Lin for helpful discussion related to this work. We thank Farshid Jafarpour for offering helpful feedback on this manuscript and Yuichi Wakamoto for sharing the experimental data. We acknowledge funding support from National Science Foundation grant DMR-1610737 (JK,EL), MRSEC at Brandeis University DMR-1420382 (JK), the Simons Foundation (JK), MRSEC at Harvard University under grant DMR-1420570 (EL,AA) and NSF CAREER 1752024 (EL,AA).

## Methods

### Cell-size regulation model

In our simulations we use an established model of cell-size regulation that has proven to accurately describe microbial growth across a range of organisms [20]. The fundamental assumption of this model is that cells grow exponentially at the single cell level and divide upon reaching a size *v*_div_(*v*_birth_) depending on their size at birth, *v*_birth_. We will assume that cells divide symmetrically so that *v*_birth_ is obtained by dividing the cell’s mother’s size at division by 2. Phenotypic variability is introduced by adding noise to both the single-cell growth rates as well as the volume at division. To this end, we take the growth rate *λ* of a cell to obey

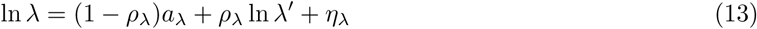

where *a*_*λ*_ is the average log growth-rate along a lineage, *ρ*_*λ*_ can be shown to approximate the Pearson correlation coefficient between mother and daughter cells, *λ*′ is the mother cell’s growth rate, and *η*_*λ*_ is a normally distributed random variable. By taking ln *λ* to progress in time in this manner, we are assuming the daughter cell’s growth rate depends only on its most recent ancestor (the mother). This will lead to a log-normal distribution of growth rates [37, 20]. Within this model, the generation time of a cell is

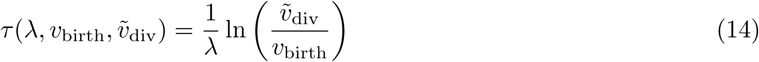

where 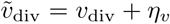 and *η*_*v*_ is another normally distributed random variable. The specific form of *v*_div_ is usually taken to be a linear combination of a size increment *v*_0_ and the size at birth,

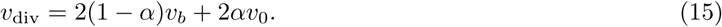

Here, the cell-size regulation strategy is controlled by the parameter *α*, for example *α* = 1 corresponds to dividing at a critical size (known as a “sizer”), while *α* = 1*/*2 corresponds to adding a constant size *v*_0_ (known as an “adder”). We refer to [1] for an in-depth discussion of the cell-size control model and its implications for population growth. Note that in the notation of our general modeling framework the phenotype that controls generation times is 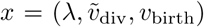 and it is straightforward to derive the distribution *f* (*x*|*x*′) from our modeling assumptions. In the special case where *α* = 1 and *η*_*v*_ = 0, the phenotype is given by *x* = ln(*λ*), the generation time is simply *τ* = ln(2)*/λ*.

It has previously been shown that the cell-size regulations strategy typically induces negative mother-daughter correlations in generation times. In the specific case where there is no growth rate variability (*η*_*λ*_ = 0), the Pearson correlation coefficient between mother and daughter cell’s generation time is −*α/*2 [20]. Growth rate and volume fluctuations tend to suppress these correlations, but preserve the sign. Therefore, since bacteria regulate their size we expect negative mother-daughter generation time correlations in the absence of any environmental noise.

### Analysis of the phenotype distributions

Here we derive the Euler-Lotka equation and present the detailed derivations of the various distributions in the finite culture.

#### Derivation of Euler-Lotka equation

Let *ψ*(*t, x, u*) denote the joint of species with phenotypes *x* and ages *u* in an instantaneous observation of a population with *N* ≫1 bacteria. We are mostly interested in the steady-state distribution *ψ*(*x, u*). In this section we will omit the superscript *F* because we are only concerned with the finite culture. Over an interval *dt*, the change in the number of cells with phenotypes between *x* and *x* + *dx* and ages between *u* and *u* + *du* is

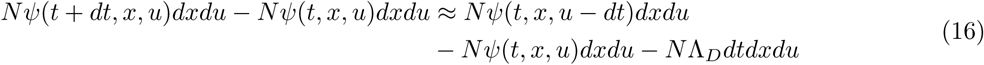

Taking the limit *dt* → 0 and dividing by *dt* gives

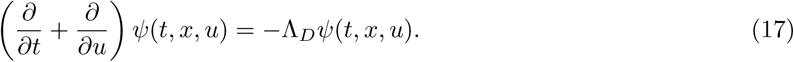

This is exactly the Von Foerster equation for an age structured population, although the role of the “death rate” is played by Λ_*D*_(*t*) and is no longer given as a model input. Instead, this must be derived from a boundary condition to ensure the population remains finite. At steady-state,

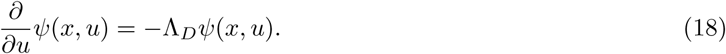

and the boundary condition is

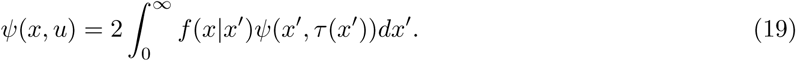

Consider the distribution of phenotypes among cells that are just born, this is given by 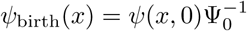 where Ψ_0_ is a normalization constant obtained by integrating 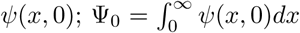. Experimentally, *ψ*_birth_(*x*) is obtained by taking an instantaneous observation of the population and sampling only the cells that have just been born. By solving Equation 18 and inserting the solution into Equation 19, it is straightforward to obtain

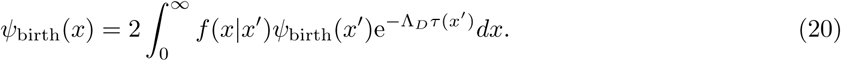

Because this distribution is time invariant, sampling those cells that have just been born at any sequence of times *t*_1_, *t*_2_, …*t*_*n*_ yields the same distribution. Carrying out this sampling procedure at an infinite sequence of times samples every cell only once, and therefore *ψ*_birth_(*x*) = *ψ*_tree_(*x*). This implies Equation 1 holds in the finite culture.

#### The current cells distribution

The distribution of phenotypes of the current cells is found by integrating *ψ*(*x, u*) over all possible ages

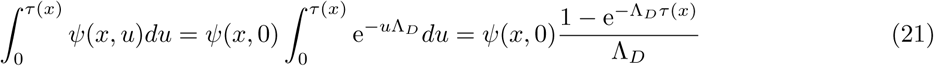

We can express the density of phenotypes among cells with age zeros as

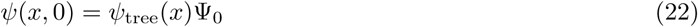

where Ψ_0_ is the prefactor in the age distribution. This gives

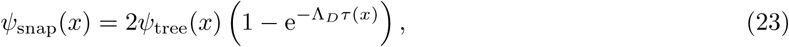

which is identical to the current cells distribution in the exponentially growing culture [30].

#### The ancestral distribution

In the finite culture *ψ*_ans_(*x*) is the distribution over all cells that have descendents in the current culture. Let 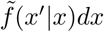 be the probability that a cell’s mother has phenotype in [*x*′, *x*′ + *dx*) given that its phenotype is *x*. Bayes’ theorem tells us that

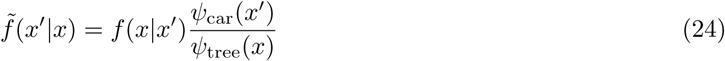

It follows that

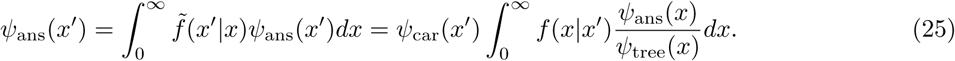

#### Division distribution

In both the continuous culture and exponentially growing culture, the carrier (or branch) distribution can be interpreted as the distribution of cells that have just divided, or the *division distribution ψ*_div_(*x*). In terms of the joint density of phenotypes and ages throughout the population, *ψ*(*x, u*), the number of cells with phenotypes between *x* and *dx* that have just divided is

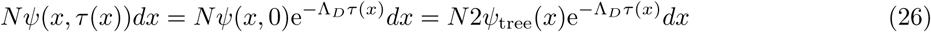

Dividing by *Ndx* gives the distribution of phenotypes among recently divided cells, which is 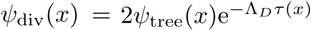. As this argument works for both a finite and exponentially growing culture, the carrier distribution, branch distribution and division distributions are all the same.

#### Summary of distributions in the finite culture

To summarize, we have discussed the following ways to sample cells in the finite culture:

- *Tree distribution* 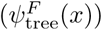: Sample all the cells throughout the history of the culture, including or excluding the current cells, but including those that are expelled before they divide.
- *Carrier distribution*) 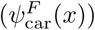: Sample all the cells throughout the culture, excluding those that are expelled before they divide.
- *Ancestral distribution* 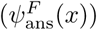: Sample all the cells in the history of the culture that have ancestors that are currently in the culture. Equivalently, select a random cell from the culture at any time and trace along a lineage backwards from this cell.
- *Current cells distribution* 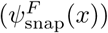: Sample all the cells that are currently in the culture.
- *Birth distribution* 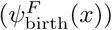: From all the cells that are in the culture at time *t*, sample only those that have just been born (meaning they have age ≈ 0), then repeat this for all *t* and average the results.
- *Division distribution* 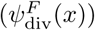: From all the cells that are in the culture at time *t*, sample only those whose age is equal to their generation time born, then repeat this for all *t* and average the results.
- The relationship between these distributions is summarized in Figure S.1.

### Derivation of Equation 4

Here we provide a more systematic derivation of Equation 4. Let *n*_*i*_ denote the number of species *i* in a culture containing *M* species. It will always be assumed that the culture is obtained by mixing homogenous cultures that have reached steady state. Let *f*_*i*_(*x*|*x*′) denote distribution of phenotypes of a daughter cell of species *i* conditioned on the mother cell’s, *ϕ*_*i*_ = *n*_*i*_*/N* be the fraction of species *i* and *ψ*_*i*_(*t, x, u*)*dudx* be the fraction of cells of species *i* with ages between *u* and *du* and phenotype between *x* and *dx*. When a cell of species *i* reaches age *τ*_*i*_(*t, x*) it divides, hence over a periods of time *dt*, the number of cells that divide is

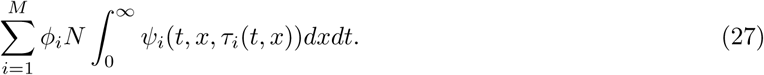

This means the per capita dilution rate is

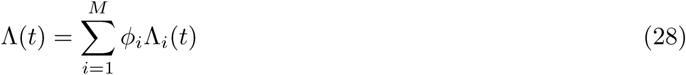

where 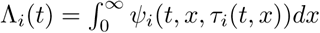 is the contributions of each species to the total dilation rate. At a given time, the number of bacteria of species *i* is *ϕ*_*i*_(*t*)*Nψ*_*i*_(*t, x, u*)*dxdu*, so over an interval *dt*, the change in the number of bacteria of species *i* with age *u < τ*_*i*_(*x*) is

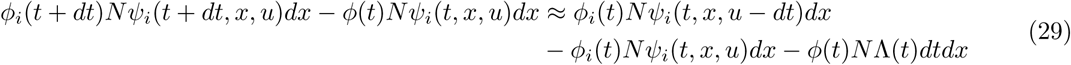

Taking the limit *dt* → 0 gives

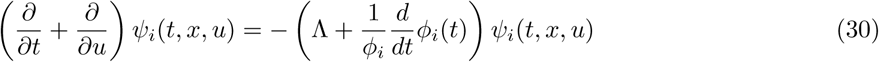

These equations depend on *ϕ*_*i*_, for which the instantaneous growth is the difference between the rate at which cells are born (Λ_*i*_(*t*)) and flushed out (Λ(*t*)). Hence,

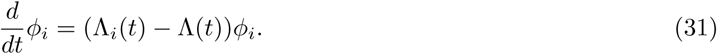

It follows that Equation 30 becomes

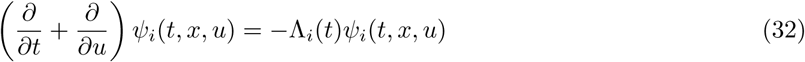

This equation holds for *u* < *τ*_*i*_(*x*) and is supplemented by the boundary conditions prescribing how the phenotypes are selected at birth,

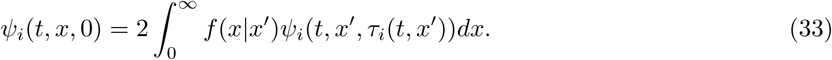

Along with an initial distribution *ψ*_*i*_(0, *x, u*), Equations 32 and 33 describe how the distribution of phenotypes and ages evolves in the mixed culture. Importantly, these equations are decoupled from *ϕ*_*i*_(*t*) and each other. Moreover, they turn out to be identical to the transport equations for a homogenous culture. This implies that the dynamics of the phenotype distribution are unchanged by the competition, and the fitness Λ_*i*_ are therefore determined by the population growth rates of the indivudual species in exponentially growing cultures. Since we have allowed for Λ_*i*_ and *τ*_*i*_ to depend on time, this result holds for both intrinsic and environmental variability.

### Interpretation of the Euler-Lotka equation in a variable environment

In order to understand how the solution of the Euler-Lotka equation relates to the true population growth rate in a temporally varying environment, we derive a generalization of the Euler-Lotka equation which can be used to bound the true population growth rate. As before, the average expected number of offspring from a randomly selected cell must be 1 in a finite culture. The expected number of offspring is twice the probability that this cell divides in the culture, thus we need to calculate this probability in order to constrain the population growth rate. To this end, let

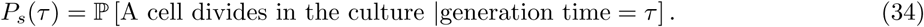

It follows that *P*_*s*_(*τ*)*ψ*_tree_(*τ*) is the joint probability that a cell has generation time *τ* and divides in the culture. Integrating this quantity yields the probability that a randomly selected cell divides in the culture, and hence the expected number of offspring from a cell is

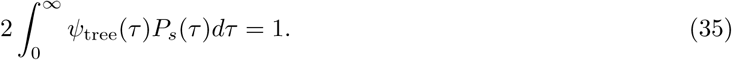

This statement assumes nothing about the underlying model of phenotypic variability, and in particular is independent of the environmental dynamics. We now introduce an explicit dependence on the state of the environment *ξ*(*t*) by defining

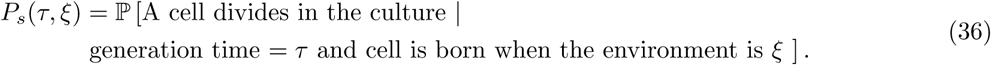

Assuming the time-scale associated with the environmental fluctuations is large relative to the the average generation time of a cell, we can make the approximation

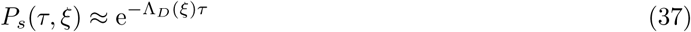

where Λ_*D*_(*ξ*) is the dilution rate in environment *ξ*(*t*) = *ξ*. Assuming the environment is a stationary process with stationary density *u*(*ξ*),

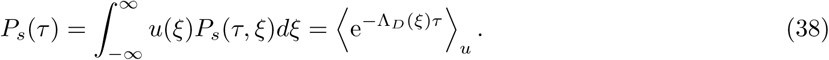

By combining Equations 35 with 38 we obtain a generalization of the Euler-Lotka equation,

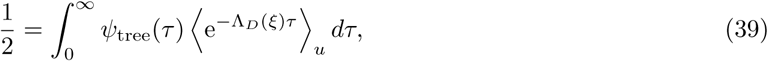

which relates the average survival probability to the tree distribution. In this context, the carrier distribution is therefore

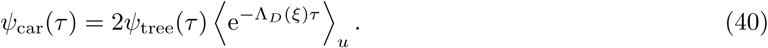

Since *ψ*_tree_(*τ*) is normalized to one, we have

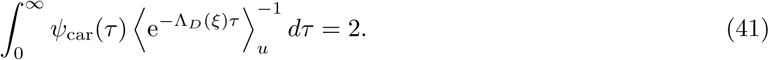

An application of Jenson’s inequality yields the bound

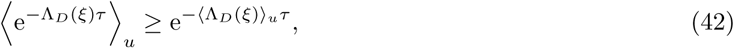

which holds for any *τ*; therefore,

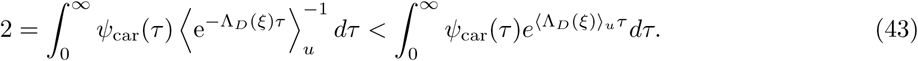

Note that Λ = ⟨Λ_*D*_(*ξ*) ⟩_*u*_ is the true population growth rate (assuming there are a large number of cells in the culture). On the other hand, since Λ_car_ satisfies Equation 6, we have

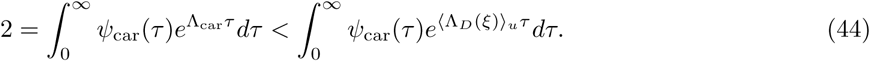

Since both integrals are decreasing functions of the exponent, this implies Λ_car_ < Λ.

We can obtain a more quantitive estimate by assuming the deviations in Λ_*D*_ from Λ_*D*_(*ξ*) and Λ_car_ are small. In this limit, Taylor expanding gives

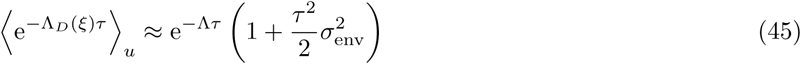

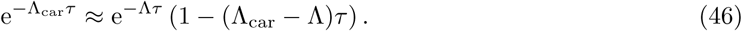

where 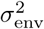 is the environmental noise term defined by 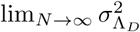. Note that as *N* tends to infinity, 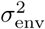 tends to 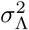, where 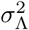 is the total variance of Λ including the contribution from intrinsic noise. Averaging Equations 45 and 46 over *ψ*_tree_(*τ*) gives

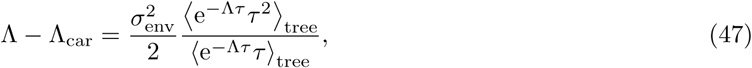

or in terms of *ψ*_car_(*τ*),

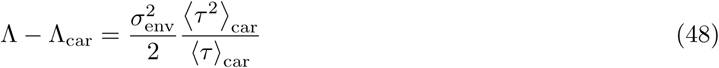

When the environmental noise is weak, Λ_*D*_ is approximately linear in the perturbation *ξ*, and therefore

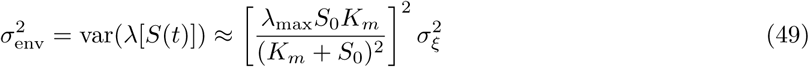

Using that

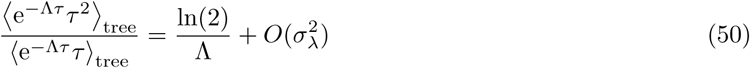

we have

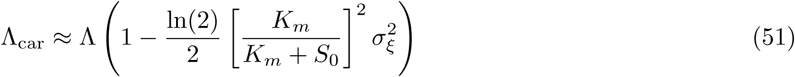

where we are using ≈ to indicate that terms of order 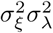 are being neglected. Note that in general *ψ*_tree_ will depend on the environmental fluctuations; however, these affects vanish in the leading order analysis. Finally, when *ξ* is modeled by an OU process, 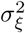 is related to the diffusion coefficient *D*_*ξ*_ and a relaxation time-scale *t*_*ξ*_ according to 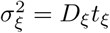.

## Supplementary figures

**Figure S.1:**
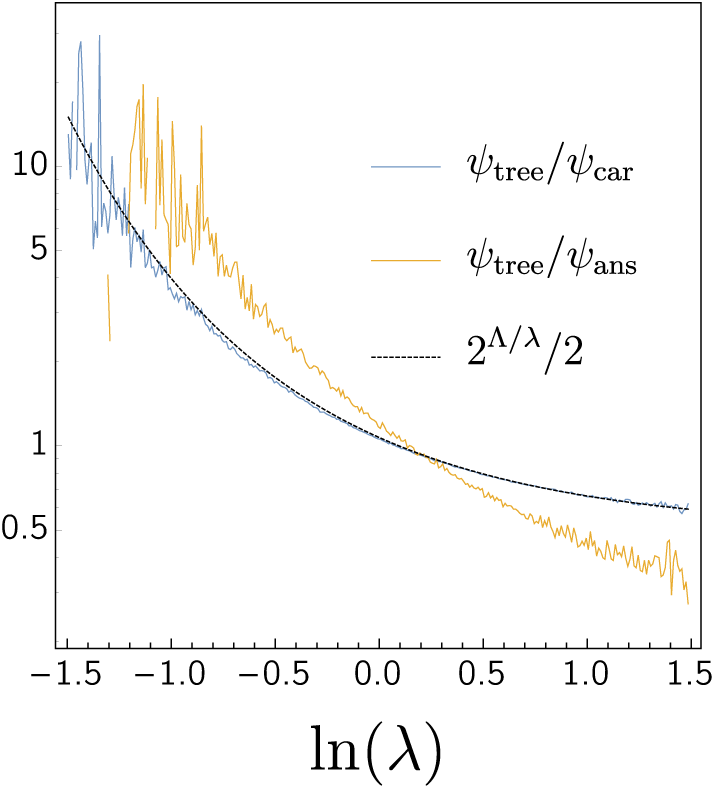
The ratio of some of the distributions plotted in Figure 4 (C). Equation 5 predicts that *ψ*_tree_(*x*)*/ ψ*_car_(*x*) = *e*^Λ*τ*(*x*)^*/*2. In this case *x* = ln(*λ*), hence 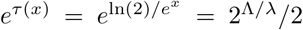. On the other hand, there is no such relationship between *ψ*_ans_ and *ψ*_tree_.

**Figure S.2:**
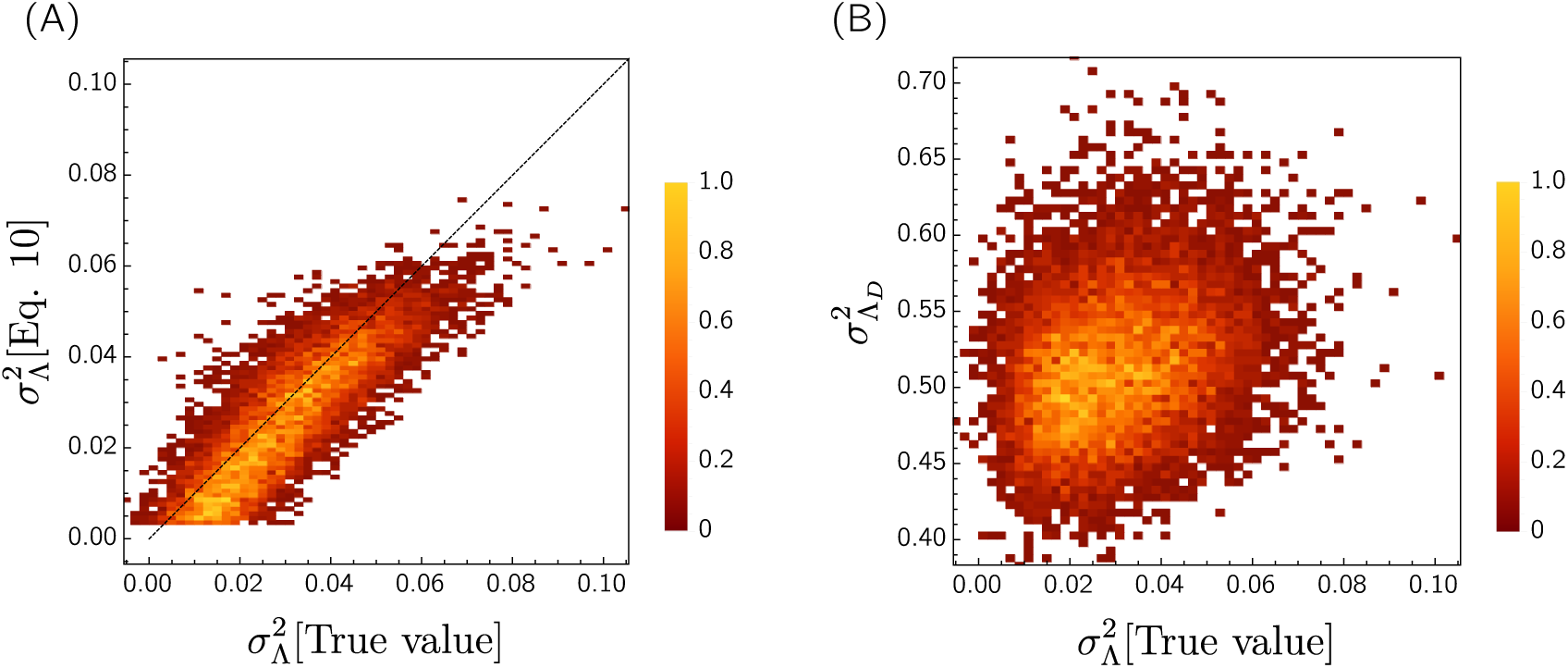
(A) Using the data plotted in Figure 5 (C), we have computed the environmental variance 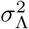 computed using Equation 10 to the true value obtained from the fluctuations in biomass accumulation rate. In order to solve for 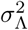 in this equation, we have computed ⟨*τ* ⟩, ⟨*τ* ^2^⟩ and Λ_in_ using the carrier distribution, while Λ is obtained by taking the average dilution rate ⟨Λ_*D*_(*t*) ⟩ and using Equation 11 to correct for population size effects. (B) The variance in the instantaneous dilution rate compared to the environmental fluctuations. This gives a much poorer estimation of the environmental variance than Equation 10.

**Figure S.3:**
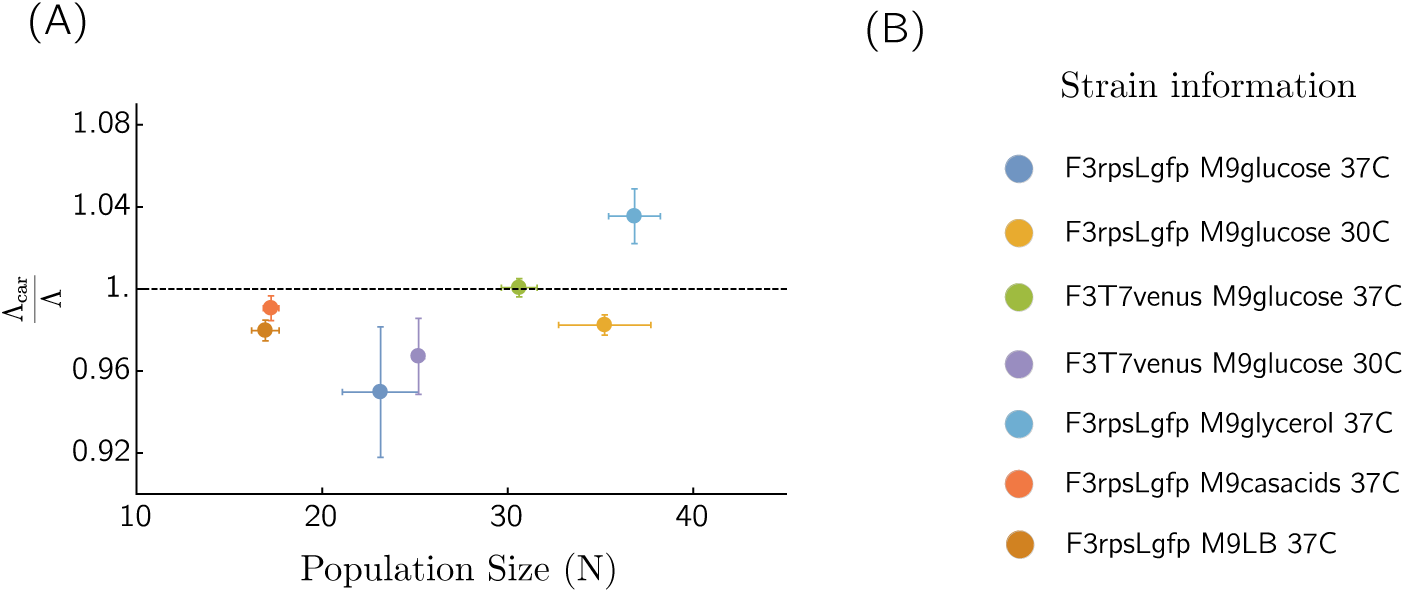
(A) The ratio Λ_car_*/*Λ obtained from different experiments as a function of the average number of cells in the culture. Each experiment corresponds to 2 - 5 biological replicates in the same experimental conditions. Λ has been obtained by computing the dilution rate and corrected for the population size. (B) Information about the different experiments, including the strain, growth medium and temperature.

**Figure S.4:**
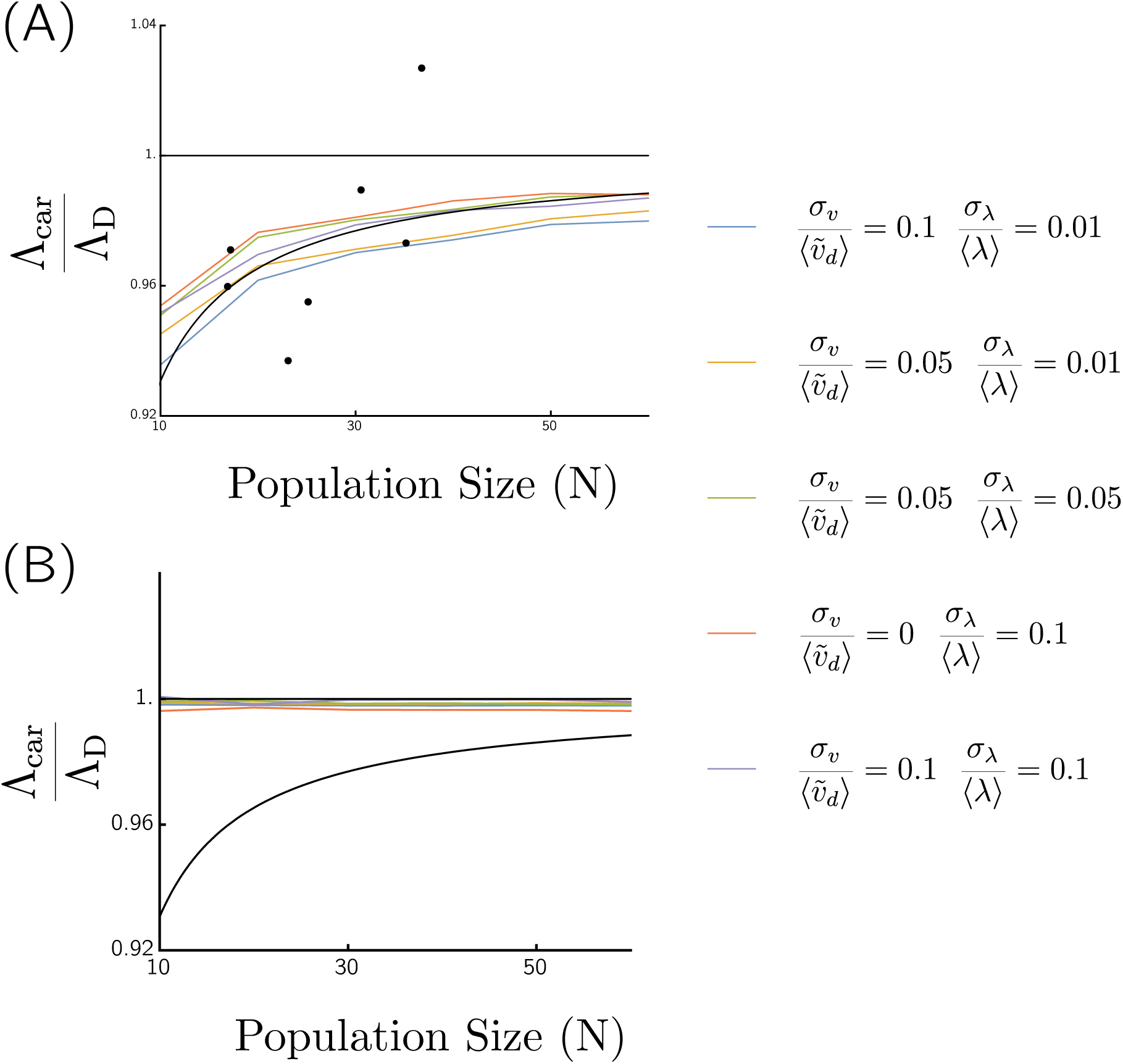
The ratios (A) Λ_car_*/*Λ_*D*_ and (B) Λ_car_*/*Λ_*D*_ for various parameters in the cell-size regulation model. The black dotes indicate the experimental data shown in detail in Figure 6 (C). Other parameters are the same as Figure 5.

## Footnotes

1 We emphasize that when computing *Ψ*_tree_(τ) one needs to know the generation times of the cells that are expelled from the culture, not the age at which they are expelled.

## References

[1] Po-Yi Ho, Jie Lin, and Ariel Amir. Modeling cell size regulation: From single-cell-level statistics to molecular mechanisms and population-level effects. Annual Review of Biophysics, 47:251–271, 2018.

[2] Avigdor Eldar and Michael B Elowitz. Functional roles for noise in genetic circuits. Nature, 467(7312):167, 2010.

[3] Zhi Wang and Jianzhi Zhang. Impact of gene expression noise on organismal fitness and the efficacy of natural selection. PNAS; Proceedings of the National Academy of Sciences, 108(16):E67–E76, 2011.

[4] Fabien Duveau, Andrea Hodgins-Davis, Brian PH Metzger, Bing Yang, Stephen Tryban, Elizabeth A Walker, Tricia Lybrook, and Patricia J Wittkopp. Fitness effects of altering gene expression noise in saccharomyces cerevisiae. eLife, 7, 2018.

[5] Michael B Elowitz, Arnold J Levine, Eric D Siggia, and Peter S Swain. Stochastic Gene Expression in a Single Cell. Science, 297(5584), 2002.

[6] Lev S Tsimring. Noise in biology. Reports on Progress in Physics, 77(2):026601, 2014.

[7] Mukund Thattai and Alexander Van Oudenaarden. Stochastic gene expression in fluctuating environments. Genetics, 167(1):523–530, 2004.

[8] Jeremie. Roux, Marc Hafner, Samuel Bandara, J. Joshua Sims, Hannah Hudson, Diana Chai, and Peter K. Sorger. Fractional killing arises from cell-to-cell variability in overcoming a caspase activity threshold. Molecular Systems Biology, 11(5):803–803, 2015.

[9] Andreas Hilfinger and Johan Paulsson. Separating intrinsic from extrinsic fluctuations in dynamic biological systems. Proceedings of the National Academy of Sciences, 108(29):12167–12172, 2011.

[10] Johan. Paulsson, Otto. G. Berg, and Mans. Ehrenberg. Stochastic focusing: Fluctuation-enhanced sensitivity of intracellular regulation. Proceedings of the National Academy of Sciences, 97(13):7148–7153, 2000.

[11] Beatrice Claudi, Petra Spröte, Anna Chirkova, Nicolas Personnic, Janine Zankl, Nura Schürmann, Alexander Schmidt, and Dirk Bumann. Phenotypic variation of salmonella in host tissues delays eradication by antimicrobial chemotherapy. Cell, 158(4):722–733, 2014.

[12] Giulia Manina, Neeraj Dhar, and John D McKinney. Stress and host immunity amplify mycobacterium tuberculosis phenotypic heterogeneity and induce nongrowing metabolically active forms. Cell host & microbe, 17(1):32–46, 2015.

[13] Yuki Sughiyama and Tetsuya J. Kobayashi. Steady-state thermodynamics for population growth in fluctuating environments. Physical Review E, 95(1), 2017.

[14] Stanislas Leibler and Edo Kussell. Individual histories and selection in heterogeneous populations. Proceedings of the National Academy of Sciences, 107(29):13183–13188, 2010.

[15] Jose M. G. Vilar and J. Miguel Rubi. Determinants of population responses to environmental fluctuations. Scientific Reports, 8(1), 2018.

[16] Mikihiro Hashimoto, Takashi Nozoe, Hidenori Nakaoka, Reiko Okura, Sayo Akiyoshi, Kunihiko Kaneko, Edo Kussell, and Yuichi Wakamoto. Noise-driven growth rate gain in clonal cellular populations. PNAS; Proceedings of the National Academy of Sciences, 113(12):3251–3256, 2016.

[17] Mats Wallden, David Fange, Ebba Gregorsson Lundius, Ö zden Baltekin, and Johan Elf. The synchronization of replication and division cycles in individual e. coli cells. Cell, 166(3):729–739, 2016.

[18] Manuel Campos, Ivan V. Surovtsev, Setsu Kato, Ahmad Paintdakhi, Bruno Beltran, Sarah E. Ebmeier, and Christine Jacobs-Wagner. A constant size extension drives bacterial cell size homeostasis. Cell, 159(6):1433–1446, Dec 2014.

[19] Guillaume Lambert and Edo Kussell. Quantifying selective pressures driving bacterial evolution using lineage analysis. Physical Review X, 5(1), 2015.

[20] Jie Lin and Ariel Amir. The effects of stochasticity at the single-cell level and cell size control on the population growth. Cell Systems, 5(4):358–367.e4, 2017.

[21] Phillip Thomas. Single-cell histories in growing populations: relating physiological variability to population growth. bioRxiv, 2017.

[22] Philipp Thomas. Making sense of snapshot data: ergodic principle for clonal cell populations. Journal of The Royal Society Interface, 14(136):20170467, 2017.

[23] Farshid Jafarpour. Cell size regulation induces sustained oscillations in the population growth rate. Physical Review Letters, 122(11), 2019.

[24] Yuichi Wakamoto, Alexander Y Grosberg, and Edo Kussell. Optimal lineage principle for age-structured populations. Evolution, 66(1):115–134, 2012.

[25] Farshid Jafarpour. Bridging the timescales of single-cell and population dynamics. Physical Review X, 8(2), 2018.

[26] Bree B. Aldridge, Marta Fernandez-Suarez, Danielle Heller, Vijay Ambravaneswaran, Daniel Irimia, Mehmet Toner, and Sarah M. Fortune. Asymmetry and aging of mycobacterial cells lead to variable growth and antibiotic susceptibility. Science, 335(6064):100–104, 2012.

[27] Lin Chao, Camilla Ulla Rang, Audrey Menegaz Proenca, and Jasper Ubirajara Chao. Asymmetrical damage partitioning in bacteria: A model for the evolution of stochasticity, determinism, and genetic assimilation. PLOS Computational Biology, 12(1):e1004700, 2016.

[28] Ariel Amir Jie Lin, Jiseon Min. Optimal segregation of proteins: Phase transitions and symmetry breaking. Physical Review Letters, 122(6), 2019.

[29] Noga Mosheiff, Bruno M.C. Martins, Sivan Pearl-Mizrahi, Alexander Grünberger, Stefan Helfrich, Irina Mihalcescu, Dietrich Kohlheyer, James C.W. Locke, Leon Glass, and Nathalie Q. Balaban. Inheritance of cell-cycle duration in the presence of periodic forcing. Physical Review X, 8(2), 2018.

[30] Jie Lin and Ariel Amir. Population growth with correlated generation times at the single-cell level. arXiv preprint arXiv:1806.02818, 2018.

[31] E. O. Powell. Growth rate and generation time of bacteria, with special reference to continuous culture. Microbiology, 15(3):492–511, 1956.

[32] Patrick Alfred Pierce Moran. Random processes in genetics. In Mathematical Proceedings of the Cambridge Philosophical Society, volume 54, pages 60–71. Cambridge University Press, 1958.

[33] David Gresham and Maitreya J Dunham. The enduring utility of continuous culturing in experimental evolution. Genomics, 104(6):399–405, 2014.

[34] Steven J. Altschuler and Lani F. Wu. Cellular heterogeneity: Do differences make a difference? Cell, 141(4):559–563, 2010.

[35] Ofer Fridman, Amir Goldberg, Irine Ronin, Noam Shoresh, and Nathalie Q Balaban. Optimization of lag time underlies antibiotic tolerance in evolved bacterial populations. Nature, 513(7518):418, 2014.

[36] Erez Lieberman, Christoph Hauert, and Martin A. Nowak. Evolutionary dynamics on graphs. Nature, 433(7023):312–316, 2005.

[37] Eric J Stewart, Richard Madden, Gregory Paul, and François Taddei. Aging and death in an organism that reproduces by morphologically symmetric division. PLoS Biology, 3(2):e45, 2005.

